# Sox2-dependent maintenance of mouse oligodendroglioma involves the Sox2-mediated downregulation of Cdkn2b, Ebf1, Zfp423 and Hey2

**DOI:** 10.1101/2020.03.04.976811

**Authors:** Cristiana Barone, Mariachiara Buccarelli, Francesco Alessandrini, Miriam Pagin, Laura Rigoldi, Irene Sambruni, Rebecca Favaro, Sergio Ottolenghi, Roberto Pallini, Lucia Ricci-Vitiani, Paolo Malatesta, Silvia K. Nicolis

**Author notes:** Northwestern University, Feinberg School of Medicine, Chicago (IL), USA. Humanitas University, Pieve Emanuele (MI), Dermatology Unit, Humanitas Research Hospital, IRCCS, Rozzano (MI), Italy.

## Abstract

Cancer stem cells (CSC) are essential for tumorigenesis. The transcription factor Sox2 is overexpressed in brain tumors. In gliomas, Sox2 is essential to maintain CSC. In mouse high-grade glioma pHGG, Sox2 deletion causes cell proliferation arrest and inability to reform tumors *in vivo;* 134 genes are significantly derepressed. To identify genes mediating the effects of Sox2 deletion, we overexpressed into pHGG cells nine among the most derepressed genes, and identified four genes, Cdkn2b, Ebf1, Zfp423 and Hey2, that strongly reduced cell proliferation *in vitro* and brain tumorigenesis *in vivo*. CRISPR/Cas9 mutagenesis, or pharmacological inactivation, of each of these genes, individually, showed that their activity is essential for the proliferation arrest caused by Sox2 deletion. These Sox2-inhibited antioncogenes also inhibited clonogenicity in primary human glioblastoma-derived cancer stem-like cell lines. These experiments identify critical anti-oncogenic factors whose inhibition by Sox2 is involved in CSC maintenance, defining new potential therapeutic targets for gliomas.

**Table of Contents Image:** 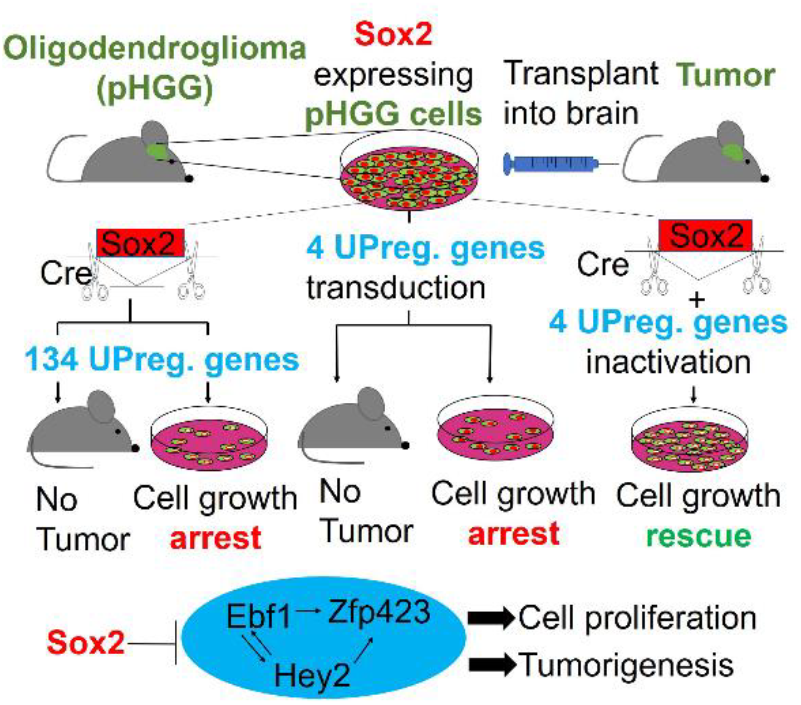

**Main Points:** Sox2 maintains glioma tumorigenicity by repressing the antioncogenic activity of a regulatory network involving the Ebf1, Hey2, Cdkn2b and Zfp423 genes.

Mutation of these genes prevents the cell proliferation arrest of Sox2-deleted glioma cells.

## Introduction

The Sox2 transcription factor has become widely known for its prominent roles in stem cells. It is important to maintain embryonic (ES), as well as various types of tissue-specific stem cells (Arnold et al., 2011; Avilion et al., 2003; Kondoh H, 2016). In the normal nervous system, Sox2 is important to maintain neural stem cells (NSC), in long-term *in vitro* culture and *in vivo*, in the hippocampus (J. Bertolini, Mercurio, S., Favaro, R., Mariani, J., Ottolenghi, S., Nicolis, S.K., 2016; J. A. Bertolini et al., 2019; Favaro et al., 2009).

Sox2 is also overexpressed in a variety of tumors (Barone, 2018; Cesarini et al., 2017; Schaefer et al., 2017). The discovery of cancer stem cells (CSC), a subset of tumor cells able to behave as “tumor-reinitiating cells”, capable to reform the tumor of origin following conventional chemotherapy (to which CSC are usually resistant) (Nicolis, 2007; Singh et al., 2004), led to the hypothesis that Sox2 may retain, in these pathological stem cells, an essential role (Barone, 2018; Nicolis, 2007; Wuebben & Rizzino, 2017).

In neural tumors, Sox2 is often active at high levels, although it is typically not mutated. This raised the question whether Sox2 is required by CSC for tumorigenesis, or whether Sox2 expression just parallels the oncogenic transformation, but has no role in maintaining transformed cells. In previous work (Favaro et al., 2014), we found that Sox2 conditional deletion in a mouse model of high-grade oligodendroglioma, obtained by PDGF-B viral overexpression (mouse PDGF-induced high-grade glioma, pHGG), completely prevents tumor reinitiation following transplantation into a mouse host brain (the assay that identifies cancer stem cells as tumor-reinitiating cells). *In vitro*, Sox2-deletion in pHGG cells significantly decreases their proliferation, and activates oligodendrocytic-like differentiation (Favaro et al., 2014). These findings paralleled those obtained in human patient-derived glioblastoma multiforme (GBM) cells, where Sox2 downregulation incapacitated cell proliferation and tumor reinitiation following transplantation (Bulstrode et al., 2017; Gangemi et al., 2009). Additionally, the importance of SOX2 in tumorigenesis and CSC was widened to various different tumor types: for example, SOX2 amplification was identified as an oncogenic driver of esophageal and lung squamous cell carcinoma, a common type of lung cancer (Bass et al., 2009), and SOX2 was found to be required within skin tumors stem cells (Boumahdi et al., 2014). An exception to these findings is represented by the Sox2-expressing neural crest-derived tumor melanoma; indeed, Sox2 conditional deletion in two different mouse models showed that Sox2 is entirely dispensable for tumorigenesis in both cases (Barone, 2018; Cesarini et al., 2017; Schaefer et al., 2017). Overall, these findings extend the significance of the study of the “dark side of Sox2” (Wuebben & Rizzino, 2017), i.e. its ability to sustain tumorigenesis, to a broad sample of tumor types.

A central remaining open question is what are the mechanisms whereby Sox2 controls CSC maintenance, i.e. which downstream target genes of Sox2 mediate its function in CSC maintenance and tumorigenesis. Previously (Favaro et al., 2014), we identified genes that are differentially expressed following Sox2 deletion in pHGG oligodendroglioma cells, and found that most of these genes are upregulated following Sox2 loss. In the present work, we individually overexpressed, within cells carrying intact Sox2, nine among the genes most upregulated following Sox2 deletion, and asked if this would reproduce, at least in part, the effect of Sox2 deletion. We identified a subset of four Sox2 downstream target genes, whose experimental overexpression (in Sox2 non-deleted cells) leads to decrease of cell proliferation and to increased cell differentiation *in vitro*, as well as to loss of tumor-initiating ability *in vivo*, pointing to these genes as important mediators of Sox2 function.

## Results

In previous work, we demonstrated an essential role for Sox2 in the maintenance of mouse PDGF-induced high-grade glioma (pHGG) stem cells, and identified genes deregulated following Sox2 deletion in these cells, representing candidate mediators of Sox2 function in pHGG stem cell maintenance (Favaro et al. 2014). In the present work, we seeked to identify, among these targets, genes that are causally involved in mediating the effects of Sox2 loss in pHGG cells: decreased cell proliferation and increased cell differentiation *in vitro*, and loss of tumor-initiating ability *in vivo* (Favaro et al, 2014).

Following Sox2 Cre-mediated deletion, the first significant changes in gene expression are detected at 96 hours after Cre transduction and consist mainly in gene upregulation, involving 134 genes (putative antioncogenes) (Favaro et al., 2014). In this work, we focused on these genes asking whether we could identify, among them, genes whose upregulation causally contributes to the antitumorigenic effect of Sox2 loss.

### Lentiviral expression in pHGG cells of a specific subset of genes upregulated following Sox2 deletion reproduces the effect of Sox2 deletion: reduced cell proliferation and increased cell differentiation

To test the effect of upregulating, in pHGG cells, individual genes found overexpressed in Sox2-deleted cells, we cloned the cDNAs of nine among the most significantly overexpressed genes (Table 1) into lentiviral expression vectors, and asked if the overexpression of any of them into Sox2-positive pHGG cells would reproduce the effects of Sox2 deletion (Fig. 1A). The cDNAs (Table 1) were cloned upstream to an IRES and a delta-NGF-receptor (dNGFR) marker gene, which is thus coexpressed with the cDNA; this allows to identify transduced cells by FACS analysis with antibodies recognizing dNGFR. pHGG cells were transduced with each of the cDNA-expressing vectors, or with a control empty vector; the percentage of transduced cells was close to 100% by FACS analysis, and all cDNAs were significantly overexpressed in comparison to empty vector-transduced control cells (not shown). At 96 hours after transduction, cells were counted; empty vector-transduced cells had proliferated and reached high numbers, comparable to those of non-transduced cell controls; however, cells transduced with the cDNAs encoding Ebf1, Cdkn2b, Zfp423 and Hey2 showed substantially lower cell numbers, ranging from 20% (Hey2) to 40% (Zfp423) of cell numbers found in controls (Fig. 1B). 48 hours after transduction, cells were also analyzed by FACS for dNGFR positivity to evaluate the percentage of transduced cells expressing the lentiviral constructs and its evolution through time (at 96 hours, 10 days and 17 days post-transduction) (Fig.1C). Cells transduced with the empty vector maintained similar, high percentages from early to late stages (still representing 80-95% of total cells at day 17); in contrast, cells transduced with Cdkn2b, Ebf1, Zfp423 and Hey2 progressively reduced their relative abundance, representing only about 10% of total cells at day 17, indicative of a disadvantage caused by the overexpressed cDNAs (Fig. 1C). In one experiment, we also evaluated the percentage of cells positive for phospho-histone H3, marking cells undergoing mitosis; phospho-histone H3-positive cells ranged between 16 and 20% in controls, but were strongly reduced (to 1-6%) in cells transduced with the cDNAs (Fig. 1D).

**Table 1.**
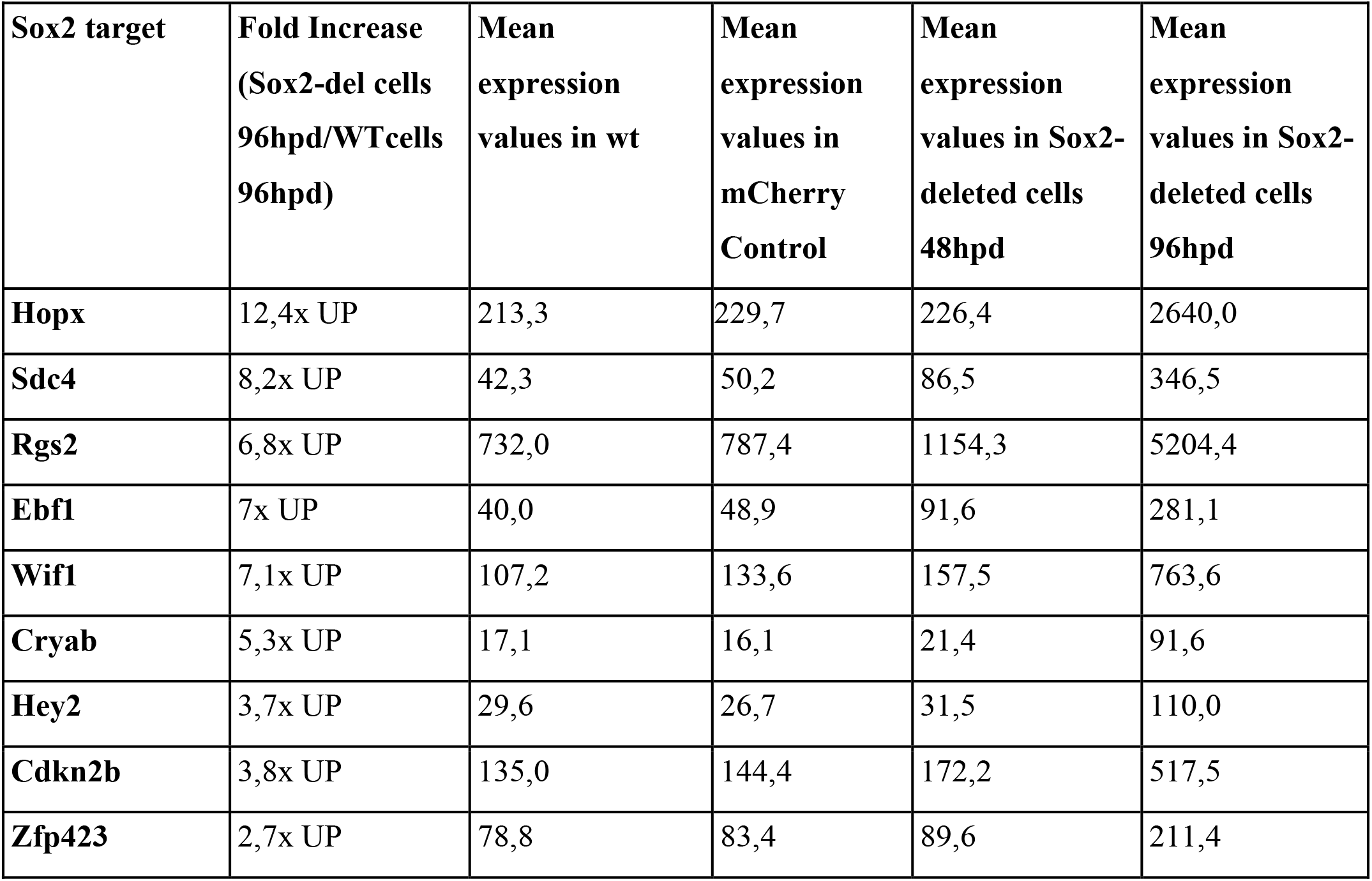
Expression of Sox2-inhibited genes in Cre-transduced Sox2-deleted pHGG cells, and in control (mCherry-transduced) cells, at 48 and 96 hours post Cre transduction hpd: hours post deletion (Expression data from Favaro et al, 2014).

**Figure 1.**
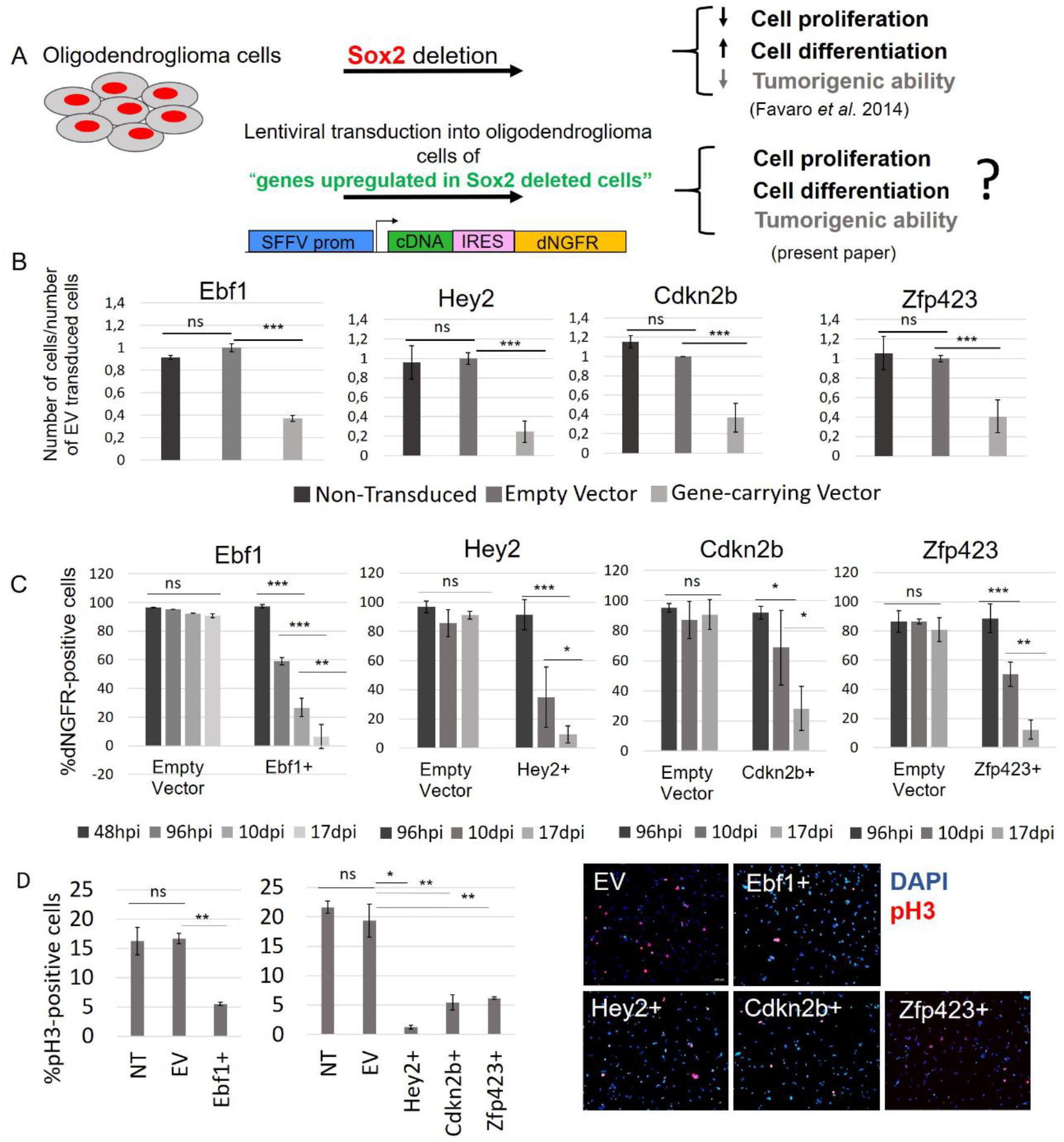
Transduction into pHGG oligodendroglioma cells of “genes upregulated following Sox2 deletion” reduces cell proliferation. A, Scheme of the rationale of the experiment, and the lentiviral vector used for upregulation of Sox2-inhibited genes. B, Cell numbers obtained following transduction of the cDNAs indicated above the histograms (“Gene-carrying Vector”), normalized to numbers obtained with Empty Vector (EV)-transduced cells, 96 hours after transduction. Error bars represent mean ± SD from at least two independent experiments performed in triplicate (***, P<0.001; Paired T test). ns: non-significant. C, Frequency (%) of dNGFR-positive (i.e. transduced) oligodendroglioma cells transduced with lentiviruses expressing the indicated cDNAs, or with Empty Vector, at different time points from the transduction, indicated below the histograms. Hpi, hours post infection; dpi, days post infection. Error bars represent mean ± SD from at least two independent experiments, performed in triplicate (***,P<0.001, **,P<0.01,*,P<0.05 Paired T test). D, Left, histograms: frequency (%) of cells undergoing mitosis (phospho-histone H3, pH3-positive) within cDNA- and Empty Vector-transduced control cells. Error bars represent mean ± SD from an experiment performed in triplicate. (***,P<0.001, **,P<0.01,*,P<0.05 Paired T test). Right: Immunofluorescence for pH3 (red) of cells transduced with empty-vector (EV), or with the indicated cDNAs.

We also asked if overexpression of these cDNAs caused cell differentiation, as previously observed for Sox2-deleted cells (Favaro et al., 2014). We analyzed cells for oligodendrocyte (GALC, O4) and astrocyte (GFAP) markers expression 7 days after transduction, by immunofluorescence (IF). Cdkn2b, Ebf1, and Zfp423 overexpression caused a significant increase (1/2%-7%) in the numbers of cells positive for oligodendrocytic (O4, GalC) and, more variably, astrocytic (GFAP) markers (Fig. 2A), with some cells exhibiting a “differentiated” morphology (Fig. 2B). This may actually be an underestimate of the number of differentiated cells induced by cDNA overexpression, because differentiation is first overt (by IF) at day 7, when the percentage of dNGFR-positive (transduced) cells is already reduced (Fig. 1C); indeed, for Zfp423, where we could evaluate the percentage of differentiated cells specifically among the transduced cells (thanks to a FLAG tag carried by the Zfp423 cDNA), the percentage of GFAP-positive cells was higher among the Flag+ than among the Flag-cells (Fig. 2A IV). With Hey2, no differentiation was observed (not shown). Finally, we did not observe cell numbers reduction, nor differentiation increase, in cells overexpressing Sdc4, Cryab, Rgs2, Wif1, Hopx (not shown). We thus continued our subsequent analyses focusing on the first group of genes, hereafter termed “Sox2-inhibited antioncogenic targets”.

**Figure 2.**
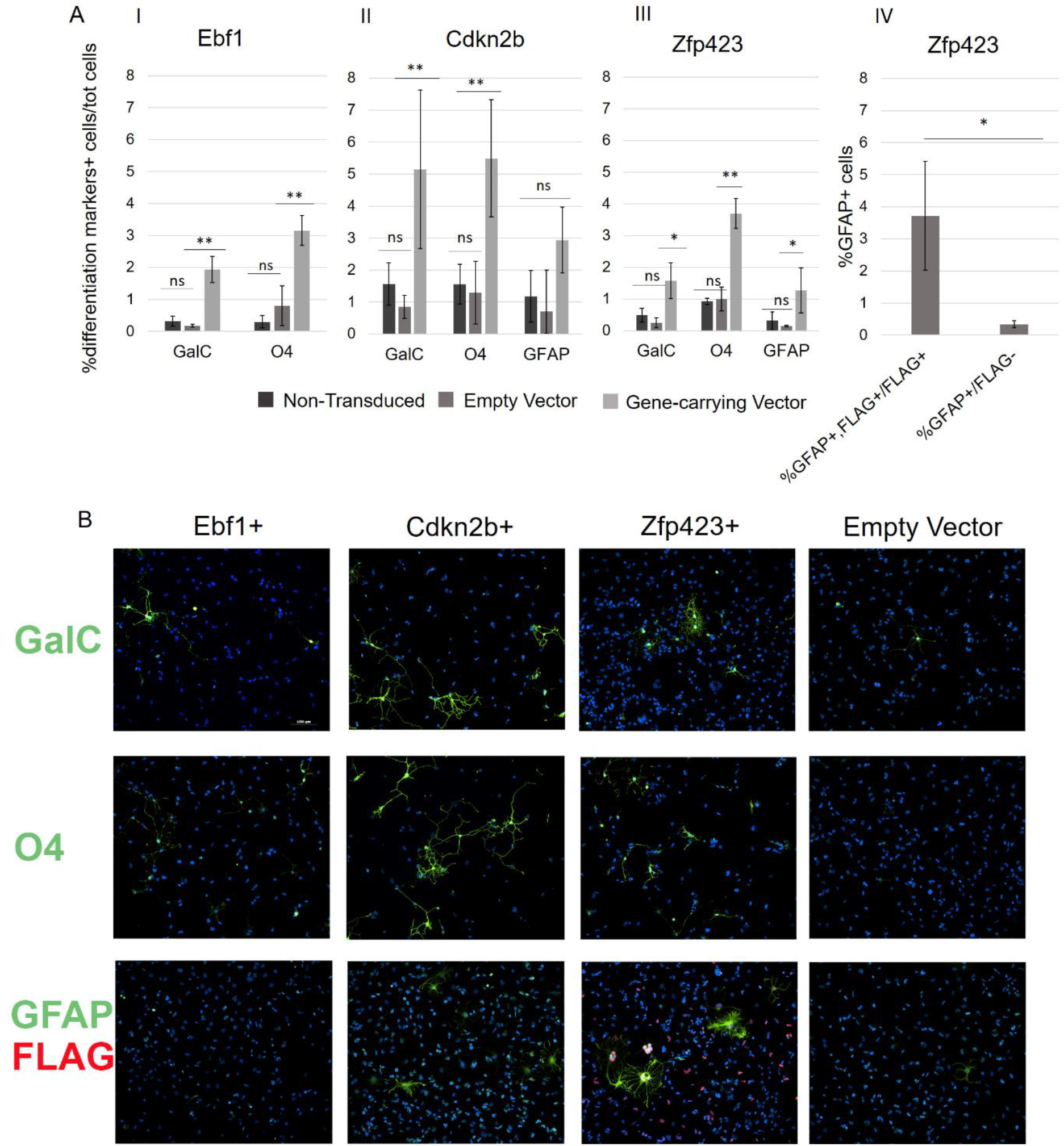
Transduction into pHGG cells of “genes upregulated following Sox2 deletion” induces glial differentiation markers and morphology in some cells. A, I-II-III: Frequency (%) of cells immunopositive for the indicated differentiation markers (GalC, O4, GFAP) within cells transduced with the cDNAs above the histograms, or with Empty Vector-transduced control cells. IV: Frequency of GFAP-positive, FLAG-positive (i.e. Zfp423-transduced) cells within total FLAG-positive cells, compared to the frequency of GFAP-positive cells within FLAG-negative cells. Error bars represent mean ± SD from at least two independent experiments, of which at least one performed in triplicate (***,P<0.001, **,P<0.01,*,P<0.05 Paired T test). B, Immunofluorescence for glial differentiation markers GalC, O4, GFAP (green) on cells transduced with the indicated cDNAs, or with Empty Vector. DAPI (blue) visualizes cell nuclei, after 7 days in culture after transduction, in serum-containing medium (scale bar, 100 um). Zfp423-overexpressing cells were double stained with antiGFAP (green) and antiFLAG (Red) antibodies.

We further asked if the Sox2-inhibited antioncogenic targets may regulate each other. Viral upregulation of Ebf1 led to significant upregulation of endogenous Zfp423 and Hey2 (5 and 3,5 fold increase, respectively, by qRT-PCR), and in turn, viral upregulation of Hey2 led to upregulation of Ebf1 and Zfp423 (Fig. 3A). Of note, Sox2 levels were unaltered in cells transduced with the four tested genes, indicating that these genes do not act on cell proliferation by reducing Sox2 expression; moreover, the changes in gene expression observed in Ebf1 and Hey2-transfected cells were not indirect effects of changes in Sox2 levels (Fig. 3B). These results point to a connection of these genes within a Sox2-dependent gene regulatory network (Fig. 3C).

**Figure 3.**
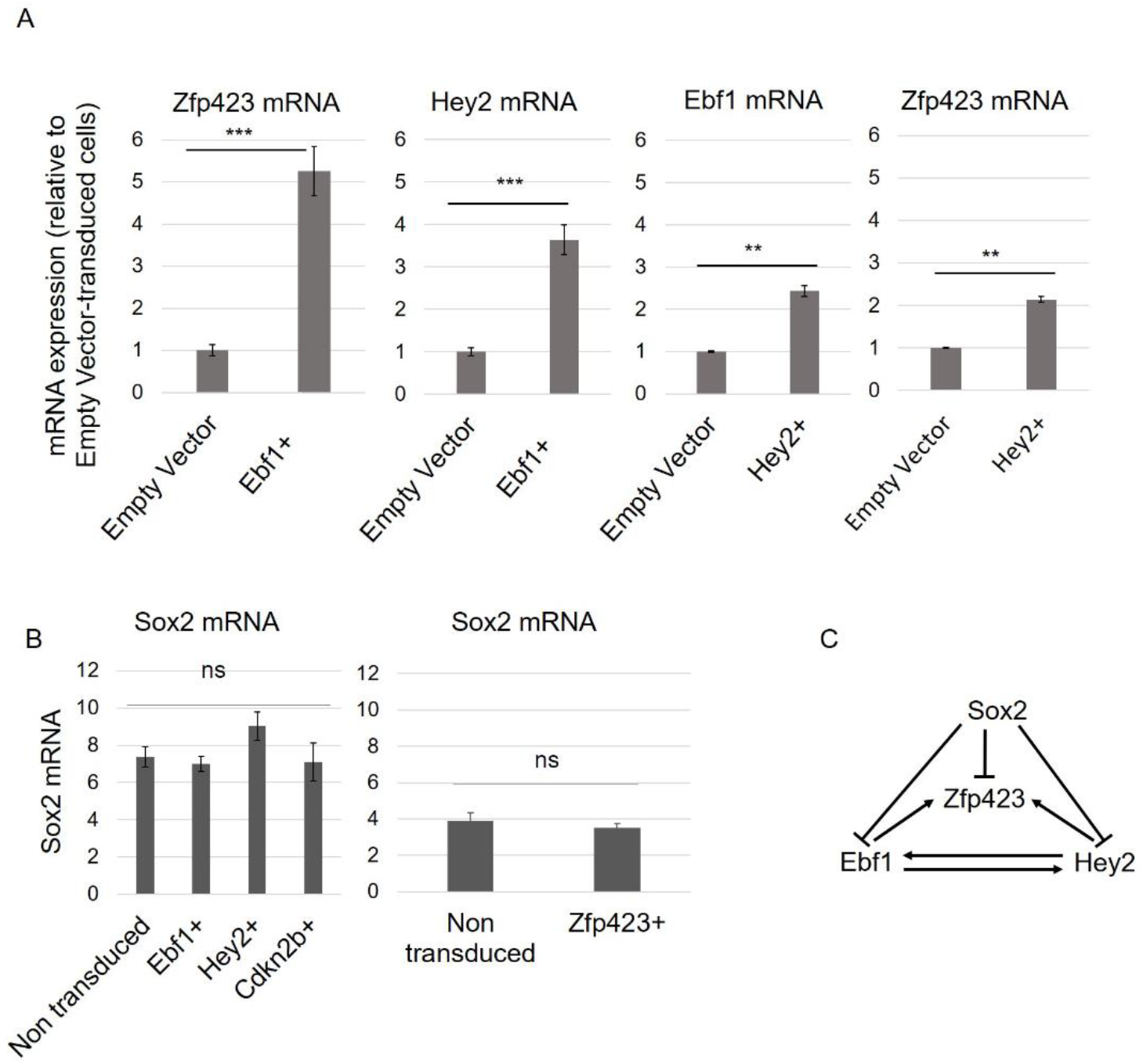
Transduction into pHGG cells of Ebf1 or Hey2 demonstrates cross-regulatory interactions between genes downstream to Sox2. A, Zfp423, Hey2 and Ebf1 mRNA (qRT-PCR) in cells transduced with Empty Vector (control, set = 1) or Ebf1 (Ebf1+) or Hey2 (Hey2+). Error bars represent mean ± SD from at least two independent experiments, performed in triplicate (***,P<0.001, **,P<0.01, Paired T test). B, Sox2 mRNA (qRT-PCR)(normalized to HPRT) in non-transduced cells or cells individually transduced with Ebf1 (Ebf1+), Hey2 (Hey2+), Cdkn2b (Cdkn2b+)(left), or Zfp423 (right). Error bars represent mean ± SD from at least two independent experiments, performed in triplicate. ns: non-significant (paired T test). C, Model for cross-regulatory interactions between Sox2, Zfp423, Ebf1 and Hey2.

### Sox2 putative antioncogenic targets act downstream to Sox2 in maintaining glioma cell proliferation

The experiments described above demonstrate anti-oncogenic/anti-proliferative activity of the 4 identified Sox2-inhibited antioncogenic targets (Cdkn2b, Ebf1, Hey2, Zfp423) upon overexpression in pHGG cells; however, is the upregulation of these genes, following Sox2 deletion, responsible for the proliferation arrest demonstrated by (Favaro et al., 2014)? To test this point, we individually mutated Ebf1, Cdkn2b or Zfp423 in pHGG cells (by CRISPR/Cas9-mediated technology), followed by Sox2 deletion by lentiviral CRE transduction (Favaro et al., 2014); we then evaluated cell numbers at 96 hours post transduction (Fig.4A). Cells where Sox2 had been deleted, and that had also been mutated in Ebf1, Cdkn2b or Zfp423 (Sox2-, Target -, Fig. 4B), showed significantly higher cell numbers, compared to cells carrying Sox2 deletion, but no mutation in the Ebf1, Cdkn2b, Zfp423 target gene (Sox2-, Target+, Fig. 4B). Importantly, ablation of each of the three target genes in cells carrying intact Sox2 genes had no significant effect, indicating that these genes importantly counteract cell proliferation only in the absence of Sox2 (when the genes are upregulated). These experiments, taken together with the upregulation of these genes shown in Sox2-deleted cells, are thus consistent with the proposal that cell proliferation arrest upon Sox2 deletion requires upregulation of the Ebf1, Cdkn2b, Zfp423 genes.

**Figure 4.**
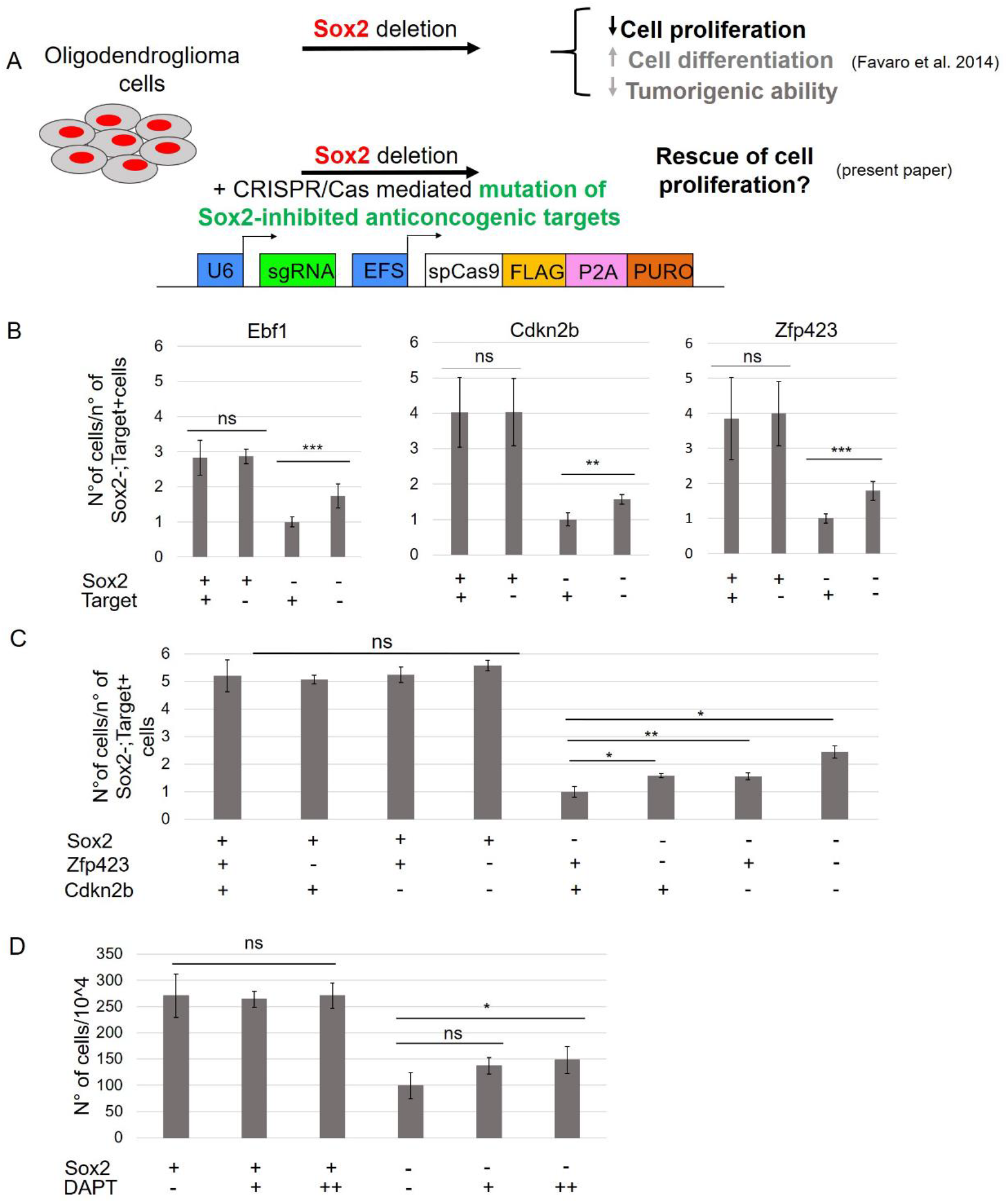
Mutation by CRISPR/Cas9, or pharmacological inhibition, of individual Sox2-inhibited antioncogenic targets partially rescues cell proliferation of Sox2-deleted oligodendroglioma cells. A, Scheme of the experiment. B, Oligodendroglioma cell numbers (counted 96 hours post Cre transduction), after CRISPR/Cas9 mutagenesis of the genes indicated above the histograms, followed (or not) by Sox2 Cre-mediated deletion. Presence of intact (+) or mutated (-) Sox2, and Cas9-targeted gene (Target +, intact; Target -, mutated), is indicated below the histograms. Cell numbers were normalized over the numbers of cells obtained in Sox2-, Target+ cells (set=1). Error bars represent mean ± SD from three independent experiments performed in triplicate (***,P<0.001, **,P<0.01, Paired T test). C, Oligodendroglioma cell numbers, following mutagenesis of both Zfp423 and Cdkn2b. Presence of intact (+) or mutated (-) Sox2, Zfp423 and Cdkn2b is indicated below the histograms. The number of Sox2-, Zfp423+, Cdkn2b+ cells is set = 1. Error bars represent mean ± SD from three experiments performed in triplicate (**,P<0.01,*,P<0.05, Paired T test). D, Oligodendroglioma cell numbers, following treatment with the Notch pathway inhibitor (DAPT) of cells carrying intact (+) or Cre-deleted (-) Sox2. Error bars represent mean ± SD from four experiments performed in triplicate (*,P<0.05, Paired T test).

We also evaluated the effect of the simultaneous mutation of two antioncogenic targets (Zfp423 and Cdkn2b), followed by Sox2 deletion; interestingly, mutating both genes had an additive effect on cell proliferation, when compared to the individual mutation of each of the genes (Fig. 4C). Finally, we evaluated the role of Hey2 in a similar type of experiment. We considered that Hey2 is downstream to the activated Notch pathway. We thus inhibited the Notch pathway by the DAPT inhibitor of the gamma-secretase enzyme, acting on the initial step of the Notch signaling pathway. The inhibitor had no effect on the proliferation of cells carrying intact copies of Sox2; however, when added in combination with Cre-mediated Sox2 deletion, it significantly increased cell numbers, as compared to cells treated only with Cre (Fig. 4D). In this experiment, Hey2 was strongly decreased by DAPT (Supplementary Fig. 1); this result is in agreement with our previous observation that Hey2 overexpression inhibits cell proliferation (Fig. 1), although it is possible that other factors downstream to Notch, for example Hes5, might also have played some role.

### Upregulation of antioncogenic Sox2 targets identified in vitro antagonizes tumorigenesis in vivo

pHGG cells carrying wild-type Sox2 quickly and efficiently give rise to tumors of the same type of the tumor of origin, following orthotopic transplantation into the mouse brain; however, Cre-mediated Sox2 deletion completely prevents tumor reinitiation (Favaro et al., 2014). To test whether the experimental upregulation of Cdkn2b, Zfp423, Ebf1, Hey2 would also be effective in antagonizing tumorigenesis in vivo, we upregulated these genes by individual transduction of the corresponding lentiviral vectors into pHGG cells, and transplanted them into host brains, to assess their tumorigenic efficiency; empty vector-transduced cells served as controls. We set up conditions to obtain a multiplicity of infection giving rise to a ratio of about 60% transduced/40% non-transduced cells (green cells and white cells in Fig. 5A). This made it possible to retrospectively analyze, in tumors, the ratio of transduced versus non-transduced cells, as a measure of the relative tumorigenic efficiency of the two cell types (Fig. 5). As shown in Fig. 5B, empty vector-transduced cells developed tumors in 5/5 transplanted brains, whereas cells transduced with the vectors upregulating the antioncogenic targets gave rise to only 1-2 tumors out of the same number of transplanted brains. The (few) tumors developing with these vectors, and those obtained with the empty vector, were then dissected, dissociated to single cells, and analyzed by FACS for their percentage of dNGFR-positive (i.e. transduced) cells. In all tumors obtained with empty vector control cells, the percentage of dNGFR-positive cells was similar to the percentage at the time of transplantation (i.e. about 60%), if not higher; however, for cells transduced with vectors overexpressing the Sox2-inhibited antioncogenic targets, the percentage of dNGFR-positive cells was importantly reduced in comparison to the percentage prior to transplantation (from about 60% to about 20-25%), indicating a disadvantage of the transduced cells in tumor formation (Fig. 5C). These findings indicate that upregulation of the Sox2-inhibited antioncogenic targets, identified in vitro, antagonizes tumorigenesis in vivo.

**Figure 5.**
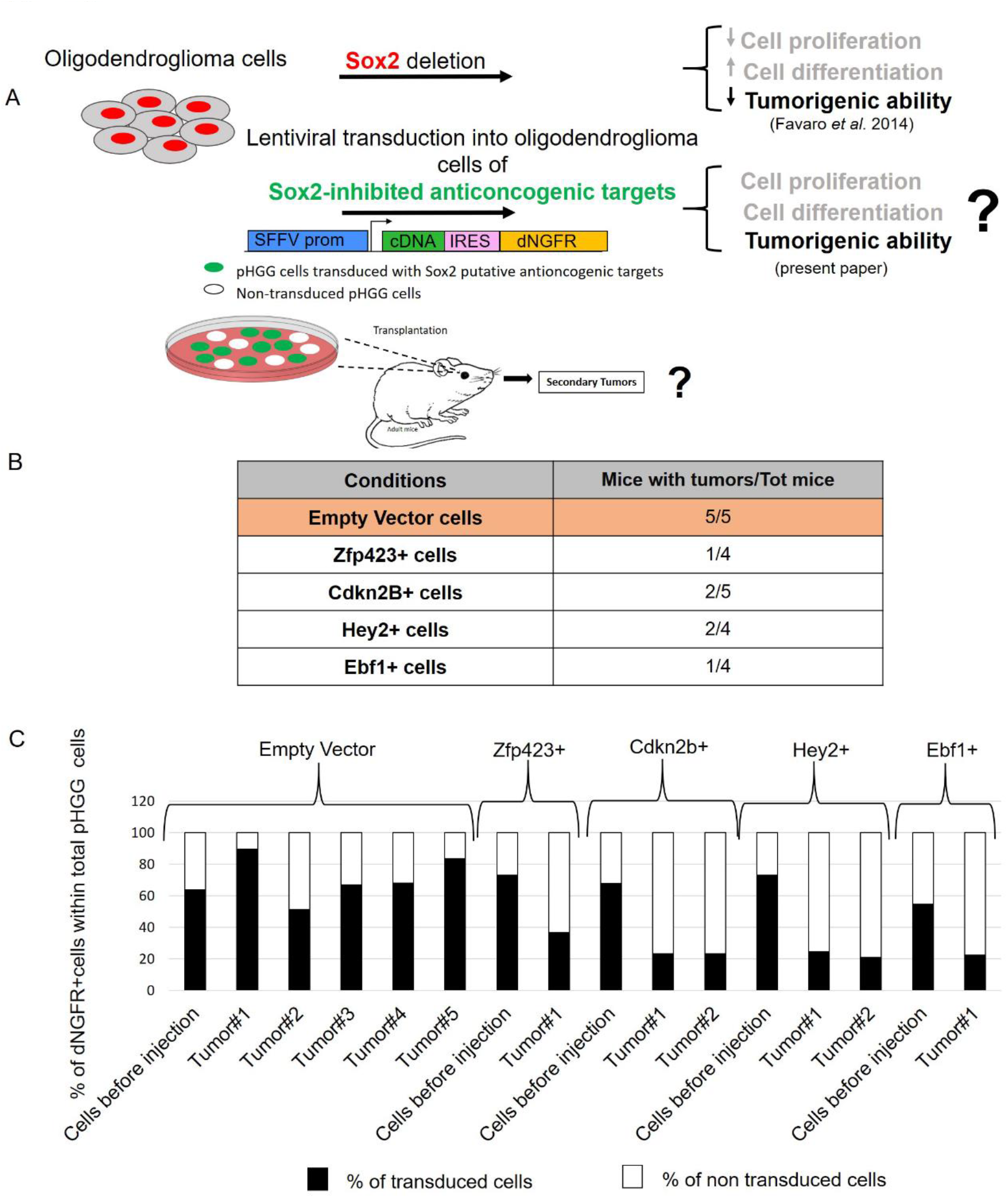
Lentiviral transduction of Sox2-inhibited antioncogenic targets antagonizes tumorigenesis *in vivo*. A, Scheme of the experiment. B, Fraction of mice developing secondary tumors following injection in the brain of cells transduced with empty vector, or the indicated cDNAs. C, Frequency (%) of dNGFR-positive (i.e. transduced) oligodendroglioma cells (black) before injection, and in different tumors obtained following injection.

### Upregulation of Sox2-inhibited antioncogenic targets identified in mouse pHGG cells antagonizes cell growth in different patient-derived, Sox2-expressing glioblastoma stem-like cell (GSC) lines

SOX2 expression has been previously documented in human glioblastoma multiforme (GBM), the most aggressive and lethal human neural tumor, and experimental downregulation of SOX2 expression in two patient-derived cell lines was shown to severely reduce tumor-reinitiating ability of the cells following transplantation into the mouse brain (Gangemi et al., 2009). We thus wished to ask if upregulation of the Sox2-inhibited antioncogenic targets, identified in pHGG cells, would also antagonize tumor cell growth of human patient-derived GSCs. We took advantage of a collection of patient-derived GSC lines (D’Alessandris et al., 2017; Marziali et al., 2016; Ricci-Vitiani et al., 2010), which all express SOX2, but at different levels (Fig. 6 and Supplementary Fig. 2). Cells from three different lines, expressing different levels of SOX2 (see Supplementary Fig. 2), were transduced with the same vectors previously used for mouse cells (encoding Cdkn2b, Zfp423, Ebf1, Hey2), and the clonogenic efficiency was tested (Fig. 6A). The cells transduced with vectors expressing the antioncogenes showed significantly reduced clonogenic efficiency, in comparison to empty vector-transduced cells, although the extent of reduction varied between different cell lines (Fig. 6B). The reduction was more pronounced for those cell lines expressing the highest SOX2 levels (GSC#1, GSC#163), as compared to the line expressing lower SOX2 (GSC#83)(Fig. 6B). These observations indicate that Sox2 targets, identified as mediators of Sox2 function in pHGG cells, also antagonize cell growth of various SOX2-expressing GBM-derived cell lines.

**Figure 6.**
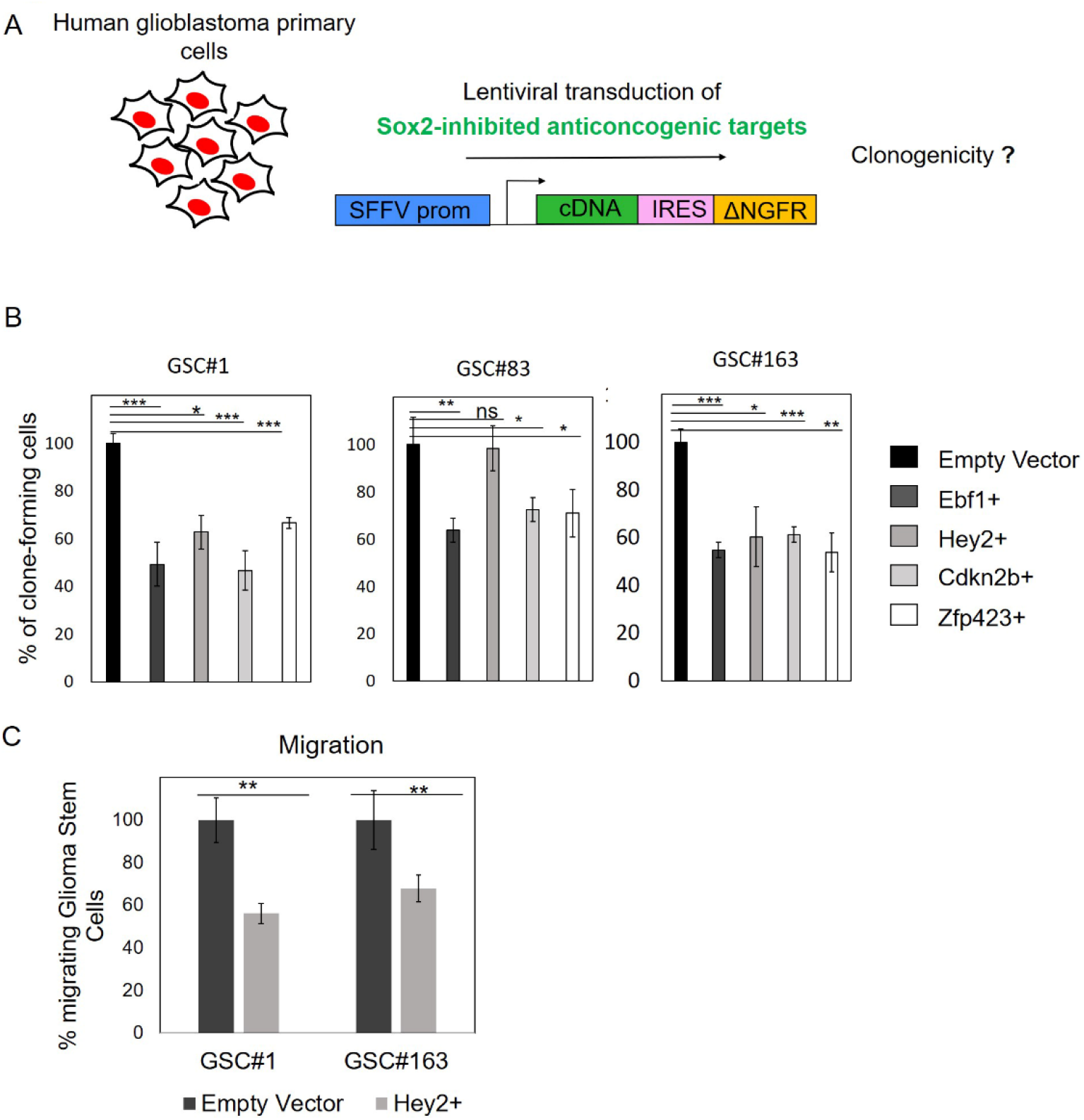
Lentiviral transduction of Sox2-inhibited antioncogenic targets reduces clonogenicity and migration in different human patient-derived glioblastoma stem-like cell lines. A, Scheme of the experiment. B, Clonogenic ability of different cell lines transduced with the cDNAs indicated on the right, or Empty Vector. GSC: glioma stem-like cells. C, Migration ability (% of migrated cells relative to Empty Vector) of cells transduced with Hey2, or Empty Vector. B,C: *, *p* <0.05; **, *p* <0.01; ***, *p* <0.001, Student’s T test. A representative experiment is shown, out of two independent experiments performed in triplicate.

Further, only Hey2 transduction (Fig. 6C), and not that of the others targets, also significantly reduced migration ability of GSC#1 and GSC#163 cells (although not of GSC#83, not shown), tested by a transwell-migration assay. Migration ability is an in vitro correlate of invasiveness, contributing to metastatic tumor development; our data suggest that Hey2 may also contribute to the regulation of this important feature of tumorigenic cells.

## Discussion

Previously, we demonstrated that Sox2 deletion arrests cell proliferation in pHGG oligodendroglioma cells and prevents in vivo tumorigenesis by such cells (a test of CSC function, Singh et al. 2008), when transplanted into mouse brain (Favaro et al. 2014). We now show that overexpression (in non-Sox2 deleted pHGG cells) of genes upregulated following Sox2 deletion inhibits their in vitro proliferation, and prevents in vivo tumorigenesis, thus mimicking the effects of Sox2 deletion.

### Sox2 maintains tumor cell properties by inhibiting four genes, able to antagonize cell growth in vitro and in vivo

In this work we identify four genes, that are importantly upregulated following Sox2 deletion in pHGG cells, as mediators of the proliferation arrest and inhibition of tumorigenesis due to Sox2 deletion. In fact, each of these genes (Cdkn2b, Ebf1, Hey2, Zfp423) significantly reduced the proliferation of non-Sox2-deleted pHGG cells, upon viral transduction (Fig. 1), and inhibited tumorigenesis after transplantation of the transduced cells in mouse brain. Not only are these genes able to reproduce the effects of Sox2 deletion in pHGG cells; their activity is also essential, as shown by CRISPR/CAS9 mutagenesis or pharmacological inhibition (Fig. 4), for repressing cell proliferation in Sox2 deleted pHGG cells. These results indicate that a critical contribution of Sox2 to the maintenance of tumorigenesis is represented by its ability to inhibit the expression of at least four genes acting as antioncogenes. Interestingly, three out of the four genes appear to be connected in a functional interaction network (Fig. 3), pointing to coordinated mechanisms for inhibiting cell proliferation stemming from Sox2 inhibition.

It is further interesting that, among the genes that we tested by overexpression, there are some, such as Hopx, Wif1 and Cryab, that are known to act as tumor suppressors in other types of tumors, although they did not affect proliferation in our experiments. It is possible that the anti-tumorigenic effect of these genes is context-dependent (i.e. specific for a given tumor type), but we cannot rule out that other types of assay might reveal a role of these genes also in pHGG cells.

Finally, initial studies of three primary human glioblastoma-derived cell lines essentially confirm a repressive role of the putative antioncogenes identified in pHGG cells, also in spontaneously arising human tumors, suggesting novel targets of possible therapeutic relevance.

### How do the identified putative antioncogenes affect tumor cell proliferation and tumorigenesis?

Of the four identified putative antioncogenes, three have previously been proposed to be somehow involved in spontaneous gliomas in man. Cdkn2b is commonly deleted in glioblastoma, often together with the adjacent Cdkn2a gene (Y. Liu et al., 2010; Melin, 2011). Ebf1 encodes a transcription factor, that is an interaction partner for TET2, an enzyme mediating DNA demethylation; TET2 is inhibited by 2-hydroxyglutarate, an oncometabolite generated by mutated isocitrate dehydrogenase (IDH) 1 and 2, which are frequently mutated in low-grade gliomas, and other tumors. As these tumors share a hypermethylated DNA phenotype, it is possible that Ebf1, as part of the TET2-Ebf1 complex, abnormally binds to the hypermethylated genes (Guilhamon et al., 2013). Furthermore, both Ebf1 and Ebf3 (a member of the family) have been found to be mutated and possibly inactivated in a variety of tumors (Liao, 2009), implying a repressive role for these genes. In particular, EBF3 is mutated in gliomas, and activates genes involved in cell cycle arrest and apoptosis while repressing genes involved in cell survival and proliferation (Liao, 2009); whether Ebf1, when increased following Sox2 deletion in pHGG cells, plays a similar role, is an interesting possibility. Hey 2, a transcription factor, is downstream to the Notch receptor signaling pathway; there are contrasting results reported in glioblastoma, regarding the effects of the activation of Notch signaling on glioblastoma cells proliferation (Giachino et al., 2015; Ying et al., 2011). Our present results are consistent with an antioncogenic effect of Hey2 activation, as proposed by Giachino et al. Finally, Zfp342 was also reported to have antigliomagenic activity, upon transfection, in one of three tested mouse astroglioma lines derived from Ink4a/Arf -/-; EGFR-mutant mice (Signaroldi et al., 2016).

From the analysis of spontaneous mutations leading to gliomas, specific pathways involving antioncogenic proteins, such as Retinoblastoma (Rb) and p53, appear to be involved, with different mutations often acting in conjunction. Thus, CDKN2A and CDKN2B are commonly deleted; they inhibit CDK4 and CDK6 kinases, which in turn activate Rb, the net effect being loss of Rb activity. Similarly, p53 may be inactivated by amplification of their inhibitor MDM proteins (Rao, Edwards, Joshi, Siu, & Riggins, 2010). In mouse, Notch signaling inactivation, combined with p53 loss, leads to the generation of aggressive brain tumors (Giachino et al., 2015); in agreement, in man Notch1 inactivating mutations are detected in gliomas, and Notch pathway effectors Hey2 and Hes5 expression levels are inversely correlated with tumor severity (Giachino et al., 2015). Our results indicate that at least two of the genes upregulated following Sox2 deletion in pHGG cells fit well in this general scheme: Cdkn2b and Hey2 increases implicate an involvement of the signaling upstream to Rb, and of at least one member of the Notch pathway. It is to be noted that the promoters of Ebf1, Cdkn2b, Zfp423 and Hey2 are directly bound by Sox2 in a human GBM-derived cell line (Fang et al., 2011). This implies, in particular for Hey2, that Sox2 might directly impact on Hey2 activity, rather than indirectly through Notch pathway modulation. A direct transcriptional repressor activity of Sox2 in neural stem cells, mediated by its interaction with the transcriptional repressor Groucho, has been reported (Y. R. Liu et al., 2014).

### Perspectives

Sox2 is overexpressed in several different tumors in man, and particularly in brain tumors. While a functional role has been demonstrated for Sox2 in the maintenance of neural tumors (Favaro et al., 2014; Gangemi et al., 2009), Sox2 mutations have not been demonstrated. It also remains unclear how Sox2 acts in the maintenance of tumors. We know from studies of wild type NSCs that, in the absence of Sox2, these cells progressively lose their replicative ability and are finally lost (Favaro et al., 2009). So far, an important mediator of Sox2 activity in sustaining long-term proliferation in vitro of NSC has been identified as Socs3, a signaling-controlling molecule (J. A. Bertolini et al., 2019). At variance with NSC, in oligodendroglioma we have now identified genes which are repressed, directly or indirectly, by Sox2, to maintain the tumor proliferation. As discussed above, these genes participate in regulatory circuits which eventually affect known antioncogenes. So, a possible mode of action of Sox2 might be to somehow repress, in gliomas, genes involved in the suppression of tumorigenesis. In the case of Cdkn2b, a gene very often deleted in GBM, Sox2 action mimics the effect of the deletion by repressing the activity of Cdkn2b.

This finding, together with observations by several authors, points to the possibility of identifying proteins which are “druggable” and thus allowing to slow down tumor progression. In this regard, a recent paper (Rubin & Sage, 2019; Walter et al., 2019) showed that mutating the Cdk2 gene in lung adenocarcinoma tumor cells sensitized these cells to the action of palbociclib, an inhibitor of Cdk4/6 kinases (themselves repressed by CDKN2b), which are frequently activated in these tumors. Our observation that Sox2 inhibits several genes which act as antioncogenes points to the exciting possibility of developing drugs able to prevent the ability of Sox2 (or interacting proteins, such as Groucho) to repress these genes.

It is also possible to propose that the products of the identified antioncogenes (e.g. the mRNA) could be delivered to tumor cells by carriers (e.g. nanoparticles) targeted to the tumor-reinitiating cells, via the recognition of specific cell surface antigens carried by them (Haas et al., 2017).

Although Sox2 itself may be envisaged as a target for therapy approaches, and immunotherapy against SOX2 significantly increased survival time in mouse models (Favaro et al., 2014), the nuclear localization of SOX2 make it a difficult target for efficient pharmacological recognition. Overall, the identification of multiple downstream Sox2 targets, impacting on various signaling pathways, representing important mediators of Sox2 function, may hopefully contribute to the design of specific, multi-hit therapy approaches.

## Materials and Methods

### Lentiviral constructs and infections

The Ebf1, Cdkn2b, Hey2 cDNAs were generated by RT-PCR from E14.5 mouse telencephalon RNA (primers: Table S 1), the FLAG-Zfp432 cDNA was cut with XhoI/AgeI from a pMSCV-puro vector (gift from G.Testa) (Signaroldi et al., 2016); all were cloned into a unique BamHI site upstream to the IRES-dNGFR cassette of the pHR SIN BX IR/EMW (Barbarani, Fugazza, Barabino, & Ronchi, 2019)(a gift from A.Ronchi, Milano).

Lentiviral vectors for CRISPR-Cas9 experiments were from Addgene: lentiCRISPRV2puro (Addgene #98290, RRID:Addgene_104990) and lentiCRISPRv2neo (Addgene #98292, RRID:Addgene_104992). The RNA guide sequences were designed and cloned according to: https://media.addgene.org/cms/filer_public/53/09/53091cde-b1ee-47ee-97cf9b3b05d290f2/lenticrisprv2-and-lentiguide-oligo-cloning-protocol.pdf, and are: Ebf1 5’-CGACAGACAGGGCCAGCCCG-3’, Cdkn2b 5’-CAGGGCGTTGGGATCTGCGC-3’, Zfp423 5’-TCACAACATCCGGCCCGGCC-3’. For Cre-encoding virus see (Favaro et al., 2014). Lentiviral particles were produced by the calcium phosphate transfection protocol in the packaging human embryonic kidney cell line 293T and infection performed as previously described (Ricci-Vitiani et al., 2004).

### Cell cultures

GSCs were isolated from surgical samples of adult patients who underwent craniotomy at the Institute of Neurosurgery, Catholic University of Rome, upon patient informed consent and approval by the local ethical committee (Pallini et al., 2008). GSC cultures were established from surgical specimens through mechanical dissociation and culturing in a serum-free medium supplemented with 20 ng/ml epidermal growth factor (EGF) and 10 ng/ml basic fibroblast growth factor (bFGF) (Peprotech, 100-15 and 100-18B) as previously described (D’Alessandris et al., 2017; Pallini et al., 2008).

### *In vitro* Sox2 target overexpression assay

For *in vitro* overexpression experiments (Fig 1–2) cells were plated at 300,000 cells/well in Matrigel-coated 6-well plates, in DMEM-F12 (Life Technologies, cod. 31331028) Complete Medium (CM), i.e. supplemented with 1ml/50ml B27 (Thermo Fisher, cod.17504044), 400μl/50ml Penicillin/Streptomicin (Euroclone, cod. ECB3001D) and 2% of Fetal Bovine Serum (FBS)(Euroclone, cod. ECS0180L). Cells were transduced 4 hours after plating, with cDNA overexpressing lentivirus or Empty Vector control lentivirus at MOI 10. Medium was changed 15 hours after transduction. After 96 hours cells were collected, counted (fig 1B), re-plated at 300.000 cells/well in Matrigel-coated 6-well plates, and collected again at 10 and 17 days post-infection. At every passage, an aliquot of cells (min 50000-max 500000) was fixed with 4% paraformaldehyde (PFA) and stained as in (Barbarani et al., 2019) with an anti-human CD271 (ΔNGFR) antibody, conjugated with Phycoerythrin (PE)(BioLegend, Cat. No. 345106, RRID: AB_2152647) (dilution 1:200) and analyzed by CytoFLEX (Backman-Coulter) to know the percentage of infected cells (Fig. 1C). To evaluate the differentiation rate, cells collected at 96 hours were plated at a density of 15,000 cells/well in Matrigel-coated 24-well plates. After 3 days, cells were fixed with 4% paraformaldehyde. To estimate proliferation transduced cells were collected 48 hours after transduction and plated at a density of 30,000 cells per well in Matrigel-coated 24-well chambered slides. After 24 hours cells were fixed with 4% PFA. Antibodies against FLAG (1:800; Sigma-Ald. F7425 RRID: AB_439687), GFAP (1:100; Millipore MAB3402 RRID:AB_94844) and phosphohistone H3 (1:1000; Millipore 06-570 RRID:AB_310177) were used for IF performed as in (Cavallaro et al., 2008). O4 and GalC IF used anti-O4 and anti-GalC hybridomas (undiluted supernatant) as in (Favaro et al., 2014), a gift from C.Taveggia (Fig 2A-B).

For *in vitro* overexpression experiments of GSCs (Fig. 6), cells were transduced with cDNA overexpressing lentivirus or empty vector as previously described (Ricci-Vitiani et al., 2004). After 48h, the efficiency of infection was evaluated. For Sox2 targets overexpression (Fig. 6B-C), cells were stained with an anti-human CD271(ΔNGFR) PE-conjugated antibody (BioLegend, Cat. No. 345106, RRID: AB_2152647) (dilution 1:200) and analyzed by FACSCanto (Becton Dickinson). Colony-forming ability was evaluated by plating a single cell/well in 96-well plates. After 3-4 weeks, each well was examined and the number of spheres/aggregates were counted.

Migration ability was evaluated by plating the cells in Corning FluoroBlok^TM^ Multiwell Inserts System (Corning Life Sciences, REF351164), according to manufacturer’s instruction. Briefly, 3×10^3^ GSCs were added to the upper chambers in stem cell medium. Stem cell medium supplemented with growth factors (EGF and b-FGF) was used as chemoattractant in the lower chambers. The plates were incubated for 48h at 37°C, then the fluorescent dye calcein acetoxymethylester (calcein-AM, Life Technologies Corporation, cod C1430) was added to the lower chamber for 30 min. The cell viability indicator calcein-AM is a non-fluorescent, cell permeant compound that is hydrolyzed by intracellular esterases into the fluorescent anion calcein and can be used to fluorescently label viable cells before microscope observation. The number of migrated cells was evaluated by counting the cells after imaging acquisition using a fluorescence microscope.

### Quantitative reverse transcriptase PCR

Primers for the quantification of mRNAs of Sox2-inhibited antioncogenic targets are reported in Table S2.

RNA isolation and Real Time PCR were performed as described in (Barbarani et al., 2019).

### CRISPR-Cas9 assays

Cells were plated at density of 1×10^6^ cells/Matrigel-coated 100mm plate, in DMEM-F12 CM, without FBS but with 10 ng/ml EGF and bFGF (Peprotech 100-15 and 100-18). Cells were transduced, after 4 hours from plating, with 15μL of lentiCRISPRV2puro or lentiCRISPRV2neo lentivirus for each plate. Medium was changed 15 hours after transduction. 48 hours after transduction cells were selected with Puromycin 5μg/ml (Sigma-Aldrich, cod. P8833) or G418 500 μg/ml (Sigma-Aldrich, cod. A1720) for 3 days. After selection an aliquot of cells (100,000-300,000 cells) was collected to evaluate the efficiency of mutagenesis. Cells treated with 2 different sgRNA (Fig. 4C) were treated first with lentiCRISPRV2puro, then selected in Puromycin for 3 days. After selection cells were collected and re-plated at density of 1×10^6^ cells in Matrigel-coated 100mm plate, subsequently treated with lentiCRISPRV2neo, than selected with G418 for 3 days. After selection cells were collected and plated at density of 300,000 cells in Matrigel-coated 6-well. After 4 hours from plating cells were transduced with Cre-encoding virus (MOI 7). Medium was changed after 15 hours. After 96 hours cells were collected and counted (Fig.4B-C). To evaluate the percentage of residual Sox2 positive cells, cells were plated at a density of 30,000 cells/well in Matrigel-coated 24-well plates. Sox2 IF was as in (Cavallaro et al., 2008). The percentage of residue SOX2-positive cells 4 days after CRE transduction (tested by IF) was comparable among different conditions, and was routinely < 10%, and never more than 30%. To evaluate the efficiency of CRISPR-Cas9-mediated mutagenesis DNA was extracted and the genomic region surrounding the sgRNA-targeted site was amplified through PCR. Primers are in Table S 3.

The amplified DNA was cloned in pGEM®-T Easy (Promega, A1360) by transformation of TOP10 *E.coli*, and inserts from individual colonies were sequenced. The sequences from CRISPR-Cas9-treated cells were compared to wild-type sequences, by using BLAST NCBI. An indel mutation was found in 100% of cases.

Notch pathway inhibition by DAPT (Sigma, cod. D5942): cells were plated at density of 300,000 cells/well in Matrigel-coated 6-well plates and, after 4 hours from plating, were transduced (MOI 7) with the Cre-encoding virus (Favaro et al., 2014). After 15 hours, medium was replaced with medium supplemented with DAPT 10μM or 2,5μM. After 96 hours, cells were collected and counted (Fig. 4D); an aliquot was used to evaluate the Hey2 and Sox2 mRNA relative abundance (Suppl. Fig. 1) by RT-PCR using primers reported in Table S 2. Sox2 residual mRNA in Cre-treated cells was <10% compared to control (not shown).

### Transplantation of virally transduced cells into mouse brain

All *in vivo* experiments were approved by the Ethical Committee for Animal Experimentation (CSEA) of the IRCCS AOU San Martino IST, Genova, and by the Italian Ministry of Health. Animals were handled in agreement with Italian current regulations about animal use for scientific purposes (D.lvo 27/01/1992, n. 116). Cell injections in the brain parenchyma of adult mice were as described (Favaro et al., 2014; Gambini et al., 2012). Cells to be transplanted were treated with viruses at MOI 5, to obtain about 60% transduced cells. We transplanted 30 000 cells per mouse of five C57Bl/6J mice per condition. Animals were monitored daily and, when the first mouse started to show signs of neurological symptoms (after 40 days), we sacrificed all the transplanted mice. The green fluorescent protein (GFP)–positive tumor area was dissected under the fluorescence microscope, and trypsinized for 20 minutes to obtain a single-cell suspension, then analysed by FACS as described previously (Barbarani et al, 2019) to assess the percentage of infected (ΔNGFR-positive) cells (Fig. 5C).

The authors declare no potential conflicts of interest.

## Acknowledgements

This research was supported by Associazione Italiana per la Ricerca sul Cancro (AIRC) grant IG 2014 – 16016 to S.K.N. C.B. is the recipient of a DIMET (Doctorate in Molecular and Translational Medicine) PhD fellowship. M.P. is the recipient of a Dipartimenti di Eccellenza fellowship. The authors wish to thank Alessandra Boe for highly qualified technical assistance in flow cytometry of human GBM cells.

## Data availability

The data that support the findings of this study are available from the corresponding author upon reasonable request.

## References

Arnold, K., Sarkar, A., Yram, M. A., Polo, J. M., Bronson, R., Sengupta, S., … Hochedlinger, K. (2011). Sox2(+) adult stem and progenitor cells are important for tissue regeneration and survival of mice. Cell stem cell, 9(4), 317–329. doi:10.1016/j.stem.2011.09.001

Avilion, A. A., Nicolis, S. K., Pevny, L. H., Perez, L., Vivian, N., & Lovell-Badge, R. (2003). Multipotent cell lineages in early mouse development depend on SOX2 function. Genes & development, 17(1), 126–140. doi:10.1101/gad.224503

Barbarani, G., Fugazza, C., Barabino, S. M. L., & Ronchi, A. E. (2019). SOX6 blocks the proliferation of BCR-ABL1(+) and JAK2V617F(+) leukemic cells. Sci Rep, 9(1), 3388. doi:10.1038/s41598-019-39926-4

Barone, C., Pagin, M., Serra, L., Motta, L., Rigoldi, L., Giubbolini, S., Badiola-Sanga, A., Mercurio, S. and Nicolis, S.K. (2018). Sox2 functions in neural cancer stem cells: the importance of the context. Insights of Neuro-Oncology, 2(1), 18–26.

Bass, A. J., Watanabe, H., Mermel, C. H., Yu, S., Perner, S., Verhaak, R. G., … Meyerson, M. (2009). SOX2 is an amplified lineage-survival oncogene in lung and esophageal squamous cell carcinomas. Nature genetics, 41(11), 1238–1242. doi:10.1038/ng.465

Bertolini, J., Mercurio, S., Favaro, R., Mariani, J., Ottolenghi, S., Nicolis, S.K.. (2016). Sox2-dependent regulation of neural stem cells and CNS development. In H. Kondoh, Lovell-Badge, R. (Ed.), Sox2, Biology and role in development and disease: Elsevier.

Bertolini, J. A., Favaro, R., Zhu, Y., Pagin, M., Ngan, C. Y., Wong, C. H., … Wei, C. L. (2019). Mapping the Global Chromatin Connectivity Network for Sox2 Function in Neural Stem Cell Maintenance. Cell stem cell, 24(3), 462–476 e466. doi:10.1016/j.stem.2019.02.004

Boumahdi, S., Driessens, G., Lapouge, G., Rorive, S., Nassar, D., Le Mercier, M., … Blanpain, C. (2014). SOX2 controls tumour initiation and cancer stem-cell functions in squamous-cell carcinoma. Nature, 511(7508), 246–250. doi:10.1038/nature13305

Bulstrode, H., Johnstone, E., Marques-Torrejon, M. A., Ferguson, K. M., Bressan, R. B., Blin, C., … Pollard, S. M. (2017). Elevated FOXG1 and SOX2 in glioblastoma enforces neural stem cell identity through transcriptional control of cell cycle and epigenetic regulators. Genes & development, 31(8), 757–773. doi:10.1101/gad.293027.116

Cavallaro, M., Mariani, J., Lancini, C., Latorre, E., Caccia, R., Gullo, F., … Nicolis, S. K. (2008). Impaired generation of mature neurons by neural stem cells from hypomorphic Sox2 mutants. Development, 135(3), 541–557. doi:10.1242/dev.010801

Cesarini, V., Guida, E., Todaro, F., Di Agostino, S., Tassinari, V., Nicolis, S., … Dolci, S. (2017). Sox2 is not required for melanomagenesis, melanoma growth and melanoma metastasis in vivo. Oncogene, 36(31), 4508–4515. doi:10.1038/onc.2017.53

D’Alessandris, Q. G., Biffoni, M., Martini, M., Runci, D., Buccarelli, M., Cenci, T., … Pallini, R. (2017). The clinical value of patient-derived glioblastoma tumorspheres in predicting treatment response. Neuro Oncol, 19(8), 1097–1108. doi:10.1093/neuonc/now304

Fang, X., Yoon, J. G., Li, L., Yu, W., Shao, J., Hua, D., … Lin, B. (2011). The SOX2 response program in glioblastoma multiforme: an integrated ChIP-seq, expression microarray, and microRNA analysis. BMC Genomics, 12, 11. doi:10.1186/1471-2164-12-11

Favaro, R., Appolloni, I., Pellegatta, S., Sanga, A. B., Pagella, P., Gambini, E., … Nicolis, S. K. (2014). Sox2 is required to maintain cancer stem cells in a mouse model of high-grade oligodendroglioma. Cancer research, 74(6), 1833–1844. doi:10.1158/0008-5472.CAN-13-1942

Favaro, R., Valotta, M., Ferri, A. L., Latorre, E., Mariani, J., Giachino, C., … Nicolis, S. K. (2009). Hippocampal development and neural stem cell maintenance require Sox2-dependent regulation of Shh. Nature neuroscience, 12(10), 1248–1256. doi:10.1038/nn.2397

Gambini, E., Reisoli, E., Appolloni, I., Gatta, V., Campadelli-Fiume, G., Menotti, L., & Malatesta, P. (2012). Replication-competent herpes simplex virus retargeted to HER2 as therapy for high-grade glioma. Mol Ther, 20(5), 994–1001. doi:10.1038/mt.2012.22

Gangemi, R. M., Griffero, F., Marubbi, D., Perera, M., Capra, M. C., Malatesta, P., … Corte, G. (2009). SOX2 silencing in glioblastoma tumor-initiating cells causes stop of proliferation and loss of tumorigenicity. Stem cells, 27(1), 40–48. doi:10.1634/stemcells.2008-0493

Giachino, C., Boulay, J. L., Ivanek, R., Alvarado, A., Tostado, C., Lugert, S., … Taylor, V. (2015). A Tumor Suppressor Function for Notch Signaling in Forebrain Tumor Subtypes. Cancer Cell, 28(6), 730–742. doi:10.1016/j.ccell.2015.10.008

Guilhamon, P., Eskandarpour, M., Halai, D., Wilson, G. A., Feber, A., Teschendorff, A. E., … Beck, S. (2013). Meta-analysis of IDH-mutant cancers identifies EBF1 as an interaction partner for TET2. Nat Commun, 4, 2166. doi:10.1038/ncomms3166

Haas, T. L., Sciuto, M. R., Brunetto, L., Valvo, C., Signore, M., Fiori, M. E., … De Maria, R. (2017). Integrin alpha7 Is a Functional Marker and Potential Therapeutic Target in Glioblastoma. Cell stem cell, 21(1), 35–50 e39. doi:10.1016/j.stem.2017.04.009

Kondoh H, L.-B. R. e. b. (2016). Sox2, Biology and Role in Development and Disease (Vol. ISBN: 978-0-12-800352-7): Elsevier, Associated Press.

Liao, D. (2009). Emerging roles of the EBF family of transcription factors in tumor suppression. Mol Cancer Res, 7(12), 1893–1901. doi:10.1158/1541-7786.MCR-09-0229

Liu, Y., Shete, S., Hosking, F., Robertson, L., Houlston, R., & Bondy, M. (2010). Genetic advances in glioma: susceptibility genes and networks. Curr Opin Genet Dev, 20(3), 239–244. doi:10.1016/j.gde.2010.02.001

Liu, Y. R., Laghari, Z. A., Novoa, C. A., Hughes, J., Webster, J. R., Goodwin, P. E., … Scotting, P. J. (2014). Sox2 acts as a transcriptional repressor in neural stem cells. BMC Neurosci, 15, 95. doi:10.1186/1471-2202-15-95

Marziali, G., Signore, M., Buccarelli, M., Grande, S., Palma, A., Biffoni, M., … Ricci-Vitiani, L. (2016). Metabolic/Proteomic Signature Defines Two Glioblastoma Subtypes With Different Clinical Outcome. Sci Rep, 6, 21557. doi:10.1038/srep21557

Melin, B. (2011). Genetic causes of glioma: new leads in the labyrinth. Curr Opin Oncol, 23(6), 643–647. doi:10.1097/CCO.0b013e32834a6f61

Nicolis, S. K. (2007). Cancer stem cells and “stemness” genes in neuro-oncology. Neurobiol Dis, 25(2), 217–229. doi:10.1016/j.nbd.2006.08.022

Pallini, R., Ricci-Vitiani, L., Banna, G. L., Signore, M., Lombardi, D., Todaro, M., … De Maria, R. (2008). Cancer stem cell analysis and clinical outcome in patients with glioblastoma multiforme. Clin Cancer Res, 14(24), 8205–8212. doi:10.1158/1078-0432.CCR-08-0644

Rao, S. K., Edwards, J., Joshi, A. D., Siu, I. M., & Riggins, G. J. (2010). A survey of glioblastoma genomic amplifications and deletions. J Neurooncol, 96(2), 169–179. doi:10.1007/s11060-009-9959-4

Ricci-Vitiani, L., Pallini, R., Biffoni, M., Todaro, M., Invernici, G., Cenci, T., … De Maria, R. (2010). Tumour vascularization via endothelial differentiation of glioblastoma stem-like cells. Nature, 468(7325), 824–828. doi:10.1038/nature09557

Ricci-Vitiani, L., Pedini, F., Mollinari, C., Condorelli, G., Bonci, D., Bez, A., … De Maria, R. (2004). Absence of caspase 8 and high expression of PED protect primitive neural cells from cell death. The Journal of experimental medicine, 200(10), 1257–1266. doi:10.1084/jem.20040921

Rubin, S. M., & Sage, J. (2019). Manipulating the tumour-suppressor protein Rb in lung cancer reveals possible drug targets. Nature, 569(7756), 343–344. doi:10.1038/d41586-019-01319-y

Schaefer, S. M., Segalada, C., Cheng, P. F., Bonalli, M., Parfejevs, V., Levesque, M. P., … Sommer, L. (2017). Sox2 is dispensable for primary melanoma and metastasis formation. Oncogene, 36(31), 4516–4524. doi:10.1038/onc.2017.55

Signaroldi, E., Laise, P., Cristofanon, S., Brancaccio, A., Reisoli, E., Atashpaz, S., … Testa, G. (2016). Polycomb dysregulation in gliomagenesis targets a Zfp423-dependent differentiation network. Nat Commun, 7, 10753. doi:10.1038/ncomms10753

Singh, S. K., Hawkins, C., Clarke, I. D., Squire, J. A., Bayani, J., Hide, T., … Dirks, P. B. (2004). Identification of human brain tumour initiating cells. Nature, 432(7015), 396–401. doi:10.1038/nature03128

Walter, D. M., Yates, T. J., Ruiz-Torres, M., Kim-Kiselak, C., Gudiel, A. A., Deshpande, C., … Feldser, D. M. (2019). RB constrains lineage fidelity and multiple stages of tumour progression and metastasis. Nature, 569(7756), 423–427. doi:10.1038/s41586-019-1172-9

Wuebben, E. L., & Rizzino, A. (2017). The dark side of SOX2: cancer - a comprehensive overview. Oncotarget, 8(27), 44917–44943. doi:10.18632/oncotarget.16570

Ying, M., Wang, S., Sang, Y., Sun, P., Lal, B., Goodwin, C. R., … Xia, S. (2011). Regulation of glioblastoma stem cells by retinoic acid: role for Notch pathway inhibition. Oncogene, 30(31), 3454–3467. doi:10.1038/onc.2011.58

