## Supplemental figures and tables for "Sox2-dependent maintenance of mouse oligodendroglioma involves the Sox2-mediated downregulation of Cdkn2b, Ebf1, Zfp423 and Hey2"

**Supporting information**

**Supplementary Fig. 1**

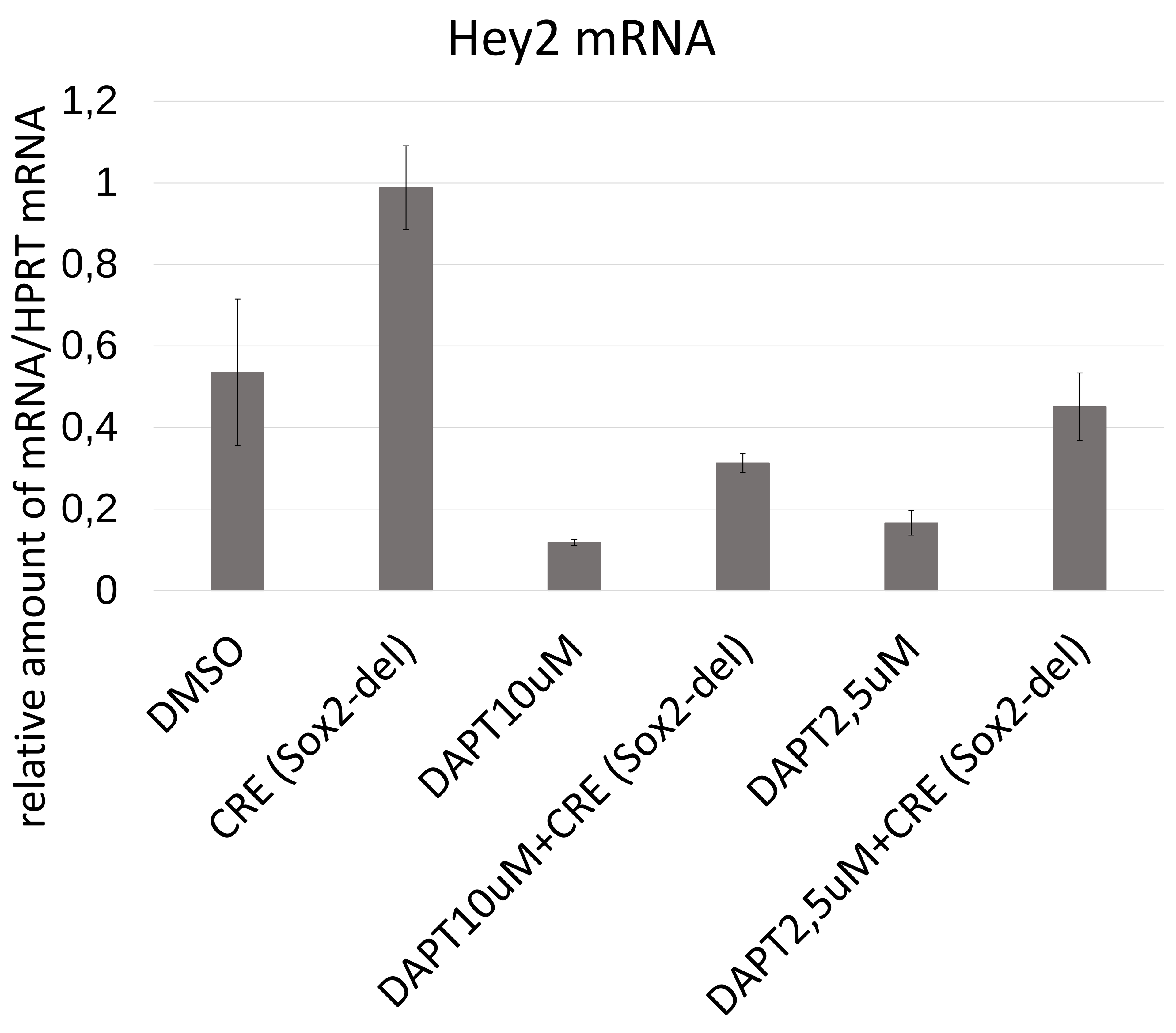


**Supplementary Fig. 1**
**Hey2 mRNA levels in DAPT-treated and untreated control cells**

**Supplementary Fig. 2**

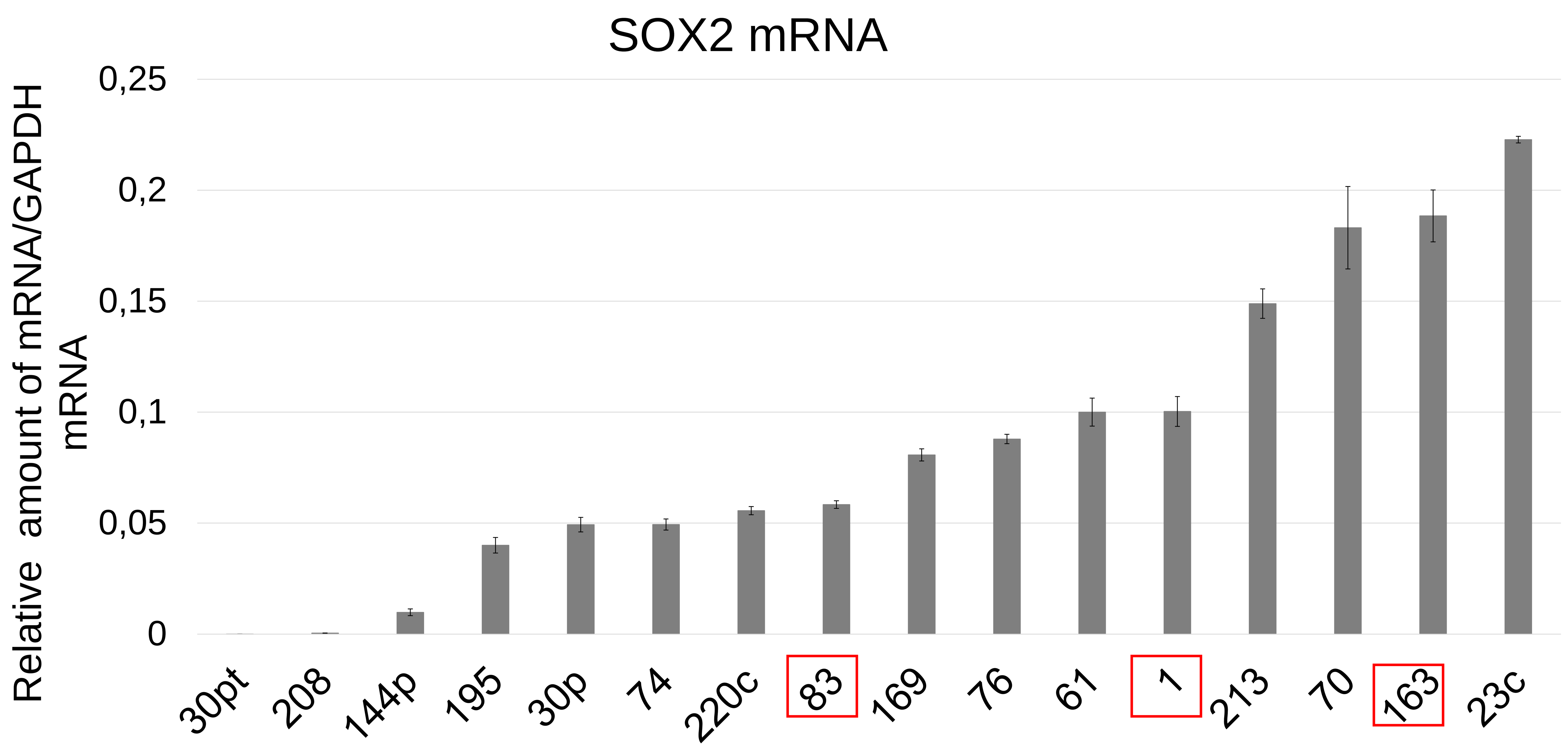


**Supplementary Fig. 2**
**SOX2 mRNA levels (relative to GAPDH) in different human primary glioblastoma cells.** Lines boxed in red are studied in Fig. 6

**Supplementary table 1**

| Gene | Forward/Reverse | Sequence |
|---|---|---|
| **Ebf1** | Forward | ACCATGGACTACAAGGACGA |
| **Ebf1** | Reverse | TCACATGGGAGGGACAATCAT |
| **Hey2** | Forward | ATATGGATCCCAGTAGCTGCTCCTCCTTCG |
| **Hey2** | Reverse | ATATGGATCCATTGCTGCTGTGTGGAACTG |
| **Cdkn2b** | Forward | ATATGGATCCCGAAGGACCATTTCTGCCAC |
| **Cdkn2b** | Reverse | ATATGGATCCTCGTGCTTGCAGTCTTCCTA |

**Supplementary table 2**

| Gene | Forward/Reverse | Sequence |
|---|---|---|
| **hEBF1** | Forward | CCTGGTGTTGTGGAAGTCACA |
| **hEBF1** | Reverse | GATGGTGGGTTCGTTGAGC |
| **mEbf1** | Forward | ACCCTGAAATGTGCCGAGTATT |
| **mEbf1** | Reverse | GGGTTTCCTGCATTCTTTAGGC |
| **mHey2** | Forward | TGGGGAGCGAGAACAATTACC |
| **mHey2** | Reverse | CCCTCTCCTTTTCTTTCTTGCC |
| **mCdkn2b** | Forward | GCCCAATCCAGGTCATGATGAT |
| **mCdkn2b** | Reverse | ATACCTCGCAATGTCACGGTG |
| **mZfp423** | Forward | ATGTCCAGGCGGAAGCAG |
| **mZfp423** | Reverse | TTTCCGATCACACTCTGGCT |
| **hSOX2** | Forward | AACCCCAAGATGCACAACTC |
| **hSOX2** | Reverse | CGGGGCCGGTATTTATAATC |
| **mSox2** | Forward | GGCAGCTACAGCATGATGCAGGAGC |
| **mSox2** | Reverse | CTGGTCATGGAGTTGTACTGCAGG |
| **hGAPDH** | Forward | ACGGATTTGGTCGTATTGGG |
| **hGAPDH** | Reverse | TGATTTTGGAGGGATCTCGC |
| **mHprt** | Forward | TCCTCCTCAGACCGGTTT |
| **mHprt** | Reverse | CCTGGTTCATTCATCGCTAATC |

**Supplementary table 3**

| Gene | Forward/Reverse | Sequence |
|---|---|---|
| **Ebf1** | Forward | GAGTGGCATTTGTCCGGTTC |
| **Ebf1** | Reverse | TTCTGAGCCCGGGACTACTA |
| **Cdkn2b** | Forward | CCAATCTAGTGCCGAGGGAT |
| **Cdkn2b** | Reverse | CTCACCGAAGCTACTGGGTC |
| **Zfp423** | Forward | TGAAGCCTAATTGCCCCTGA |
| **Zfp423** | Reverse | CCCTTGGGAAGTGGCCTATG |
